# Representation of uncertainty in macaque visual cortex

**DOI:** 10.1101/724021

**Authors:** Olivier J. Hénaff, Zoe M. Boundy-Singer, Kristof Meding, Corey M. Ziemba, Robbe L. T. Goris

**Author notes:** Correspondence should be addressed to R.L.T.G.

## Abstract

Uncertainty is intrinsic to perception. Neural circuits which process sensory information must therefore also represent the reliability of this information. How they do so is a topic of debate. We propose a view of visual cortex in which average neural response strength encodes stimulus features, while cross-neuron variability in response gain encodes the uncertainty of these features. To test our theory, we studied spiking activity of neurons in macaque V1 and V2 elicited by repeated presentations of stimuli whose uncertainty was manipulated in distinct ways. We show that gain variability of individual neurons is tuned to stimulus uncertainty, that this tuning is invariant to the source of uncertainty, and that it is specific to the features encoded by these neurons. We demonstrate that this behavior naturally arises from known gain-control mechanisms, and derive how downstream circuits can jointly decode stimulus features and their uncertainty from sensory population activity.

---

Sensory systems offer a window onto a world that cannot be known perfectly. Uncertainty about the world can arise externally, when sensory cues are incomplete or contradictory, or internally, when noise corrupts neural representations. Ideal perceptual systems take this uncertainty into account: if a sensory cue is ambiguous, prior experience guides its interpretation ^1^, and when multiple cues are available, they are combined in proportion to their reliability ^2^. When humans and other animals perform perceptual tasks, they often follow these normative predictions ^3–6^.

These behavioral effects imply that the neural circuits which mediate perception assess the uncertainty of sensory information. How they do so is unclear. A prominent hypothesis is that the same neurons that encode a stimulus feature also encode the uncertainty about this feature ^7–9^. However, which aspect of neural activity represents uncertainty remains a topic of debate. Response variability is a promising candidate: in visual cortex, it is maximal in the absence of a stimulus ^10^, and declines with contrast ^11^, aperture size ^12^, and attention ^13,14^. Since each of these factors is associated with increased information about the visual environment, response variability might represent stimulus uncertainty ^9^.

Here, we incorporate this hypothesis in the canonical model of neural coding. We propose that, while average response magnitude encodes stimulus features, variability in response gain encodes the uncertainty of these features. We formalize this proposal in a doubly stochastic response model in which spikes arise from a Poisson process whose rate is the product of a deterministic response mean and a stochastic response gain. The mean response is governed by a parametric function commonly referred to as the *classical receptive field*. We introduce a second function, the *uncertainty receptive field*, which determines the variance of the response gain.

To test our theory, we studied responses of individual orientation-selective neurons in macaque visual cortex, driven by repeated presentations of stimuli whose orientation uncertainty was manipulated in two different ways. As predicted, we found that gain variability selectively depends on stimulus uncertainty, and that this selectivity was invariant to the source of uncertainty. This appears to be a general property of visual coding: we find that the gain variability of texture-selective neurons in V2 systematically increases with an image’s textural uncertainty. To identify the neural computation that gives rise to this behavior, we developed a probabilistic model of divisive normalization in which driving input is divided by *noisy* suppressive inputs. This model quantitatively matches the effects of stimulus uncertainty on response variability.

Finally, we asked whether our coding scheme permits down-stream circuits to quickly decode the information needed for perceptual tasks. We find that neuronal gain exhibits slow dynamics, not fast. Consequently, gain variability cannot be readily decoded from individual neurons. We derived the optimal decoder of neural population activity, and used model simulations to investigate its performance. We show that stimulus orientation and gain variability can be jointly decoded from a brief V1-population response and that gain variability faithfully predicts the accuracy of orientation decoding. Together, these results establish cross-neuron variability in response gain as a candidate currency of uncertainty in sensory cortex.

## RESULTS

In primary visual cortex (V1), neurons are tuned for local image orientation, making this area well suited to inform perceptual orientation estimates. The optimal estimation strategy is to consider the probability of each possible orientation given the V1 population response, and select the value that is most likely. However, because of internal and external noise, this likelihood function and the resulting orientation estimates vary from trial to trial (Fig 1a, left). The lower the signal-to-noise ratio, the greater the uncertainty, and the greater the variance of the optimal estimate (Fig. 1a, right).

**Figure 1.**
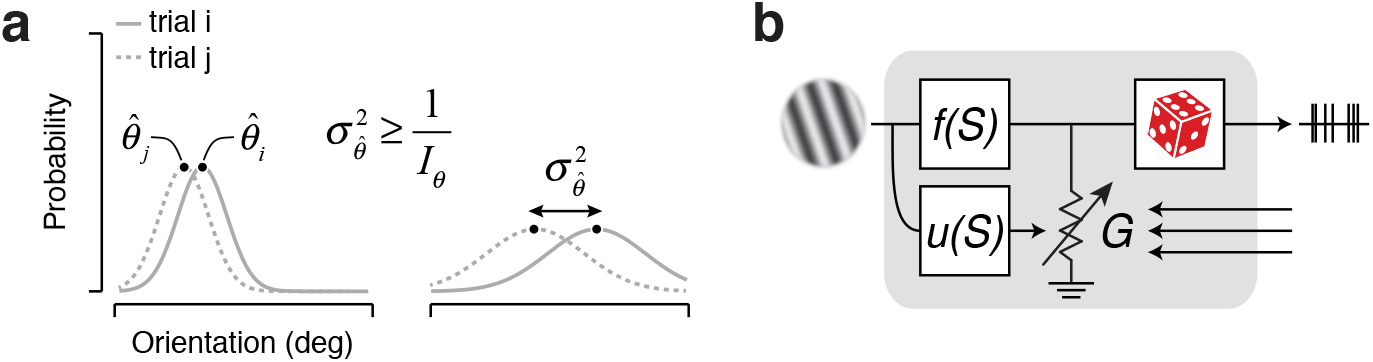
Encoding the information needed for perceptual tasks. (**a**) The optimal strategy to estimate a perceptual quantity such as local image orientation is to compute the likelihood of each possible stimulus value (grey lines) and choose the most likely option (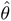 black dots). Due to noise, this value will differ across trials (full vs dotted lines, 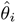 vs 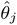). The larger the uncertainty of the sensory information, the wider the likelihood function, and the greater the variance in the perceptual estimate (left vs right panel). The variance of the optimal estimate 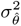 is larger than or equal to the inverse of the Fisher information *I*_*θ*_ ^20^. (**b**) Schematic summarizing the proposed model. Spikes arise from a Poisson process whose rate is the product of a deterministic drive *f* (*S*) and a stochastic gain *G*. *f* (·) governs the mapping of stimulus features onto drive and hence controls response mean; *u*(·) governs the mapping of stimulus uncertainty onto gain variability and hence controls response variance.

Many perceptual tasks require that the uncertainty of perceptual estimates be assessed on a moment-by-moment basis. How can downstream circuits instantaneously assess the reliability of V1 orientation reports? Since this reliability varies systematically with certain features of the stimulus such as the size and contrast of a local image patch, we hypothesize that V1 neurons might encode reliability through a separate channel tuned to these features. Specifically, let us assume that a neuron’s response is in part governed by a deterministic function of the stimulus *f* (*S*) (the ‘classical receptive field’) and in part by noise (Fig 1b, top branch). Previous work has shown that spiking activity is well described by a modulated Poisson process whose rate is the product of *f* (*S*) and a stochastic response gain *G* ^15^. In particular, if the gain *G* has a unit mean and varies on a time-scale which is slow relative to the measurement interval Δ*t*, spike count variance can be decomposed as:

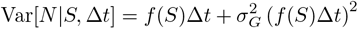

The first term is the variance due to the Poisson process, the second is due to variability in the firing rate and grows with the variance of the gain 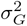. Whereas this gain variance was originally assumed to be stimulus-independent ^15^, we propose that it systematically depends on the stimulus via an ‘uncertainty receptive field’ *u*(*S*) (Fig 1b, bottom branch). If the uncertainty receptive field is selective for stimulus features that induce uncertainty, gain variability may provide a useful assay for the reliability of V1 orientation reports.

The classical receptive field is associated with two key properties: it endows sensory neurons with a particular selectivity and a particular invariance. For example, the firing rate of V1 complex cells reports the total amount of energy at a particular orientation, irrespective of the image’s polarity or precise location within the receptive field ^16^. We hypothesize that the computations underlying the uncertainty receptive field achieve a similar effect. Specifically, we expect that the gain variability of sensory neurons reports the total amount of uncertainty about the features they represent, while being invariant to the source of this uncertainty.

### Testing the theory in visual cortex

To test our theory, we analyzed responses of neurons in macaque visual cortex elicited by mixtures of sinusoidal gratings (Fig. 2a; a model-based analysis of these data concerned with mechanisms of orientation selectivity has been previously published ^17^). These stimuli are Gaussian-distributed in the orientation domain, hence the perceptual uncertainty about their orientation depends on only two factors: the total amount of stimulus energy (contrast), and its dispersion (spread). Indeed, increasing stimulus spread increases perceptual discrimination thresholds because it acts as external orientation noise ^18^. Reducing stimulus contrast has the same effect because it exposes internal noise ^19^.

**Figure 2.**
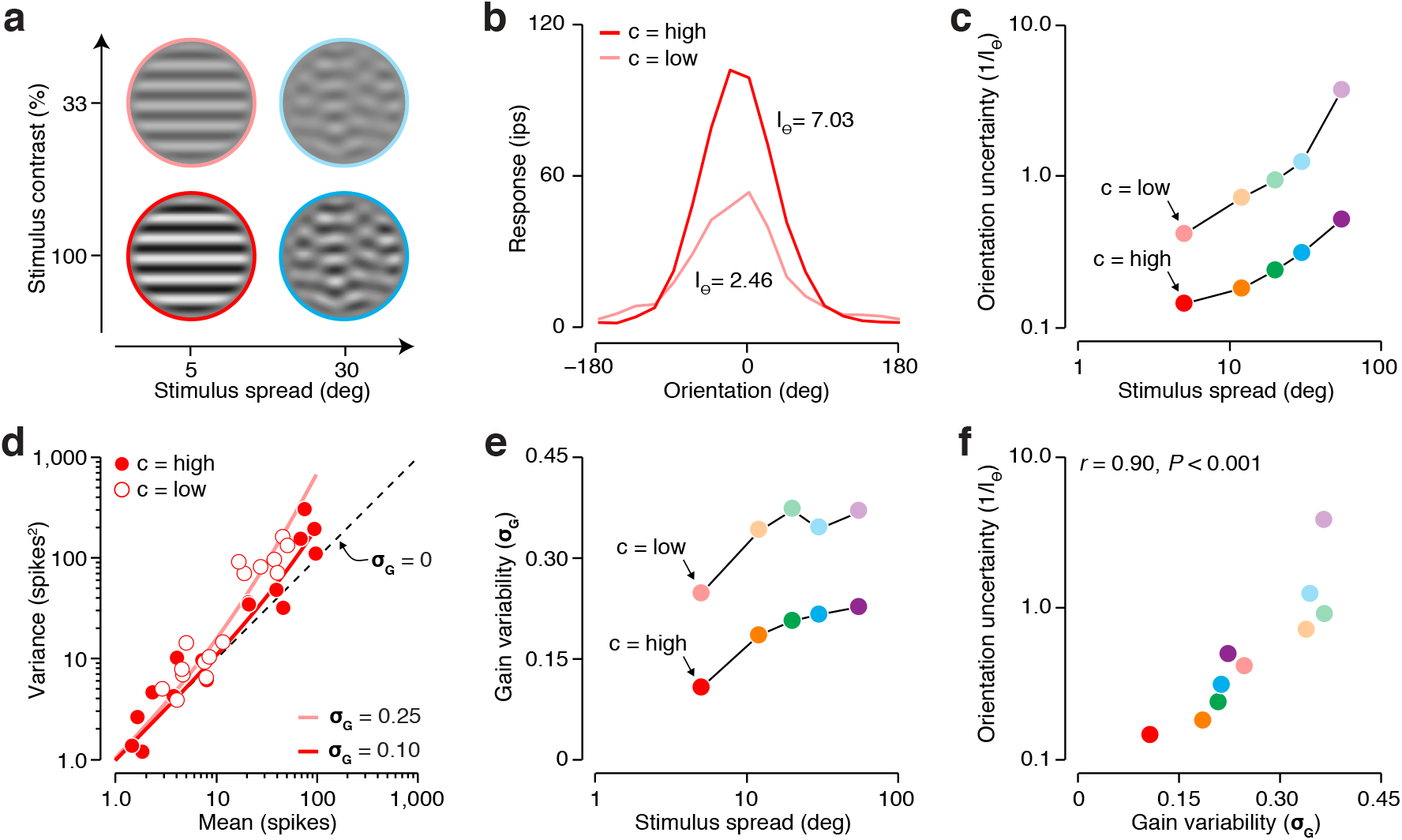
Estimating stimulus uncertainty and gain variability. (**a**) Each stimulus consisted of a sum of drifting sinusoidal gratings, with drift directions drawn from a Gaussian distribution. Stimuli differed in center drift direction (16 levels), spread (5 levels, represented by hue), and contrast (2 levels, represented by saturation). (**b**) Mean responses of a V1 neuron measured for high- and low-contrast narrow-band stimuli drifting in different directions. Responses were computed by counting spikes in a 1,000 ms window following response onset. From these tuning curves, we estimated Fisher Information (*I*_*θ*_, see online Methods). We use its inverse as a measure of orientation uncertainty. (**c**) Orientation uncertainty as a function of stimulus contrast and spread for the example neuron. (**d**) Variance-to-mean relation of the example neuron for the narrow-band stimuli. Different points indicate different drift directions. Lines illustrate the predictions of the modulated Poisson model, fit separately to the high- and low-contrast conditions. (**e**) Gain variability as a function of stimulus contrast and spread for the example neuron. (**f**) Orientation uncertainty as a function of gain variability for the example neuron.

These behavioral effects are mirrored by changes in coding capacity at the level of individual neurons. Consider the orientation information encoded in the response of an example neuron to a narrowband stimulus. Reducing stimulus contrast from 100 to 33% approximately halved this neuron’s mean response (Fig. 2b). To determine the impact of this loss of responsivity, we estimated the Fisher information associated with both conditions (*I*_*θ*_, see Online Methods). This statistic quantifies the amount of orientation information that can be extracted from the neuron’s responses by an optimal decoder. Its inverse provides a lower bound on the variance of the optimal estimate ^20^, and we use it here as a proxy for orientation uncertainty. For the high-contrast stimulus, the Fisher information was 7.03; for the low-contrast stimulus, it was 2.46 (Fig. 2b). For this neuron, the contrast reduction thus led to a substantial increase in orientation uncertainty. Increasing stimulus spread had the same effect, which was evident both at high and low contrast (Fig. 2c).

Are these changes in stimulus uncertainty reflected in the neuron’s gain variability? We used the modulated Poisson model to estimate gain variability for each stimulus family separately (Online Methods). For the narrow-band stimulus, gain variability was greater at low contrast than at high contrast (Fig. 2d; *σ*_*G*_ = 0.10 at high contrast, *σ*_*G*_ = 0.25 at low contrast). Moreover, gain variability also increased with stimulus spread, irrespective of the contrast level (Fig. 2e). Across all stimulus families, orientation uncertainty and gain variability exhibited a striking quantitative relationship (*r* = 0.90, *P* < 0.001; Fig. 2f).

The dependency of gain variability on stimulus uncertainty was evident across the population of V1 and V2 neurons. There was some heterogeneity in the effects of the stimulus manipulations on neurons’ responses, but overall, both manipulations substantially increased orientation uncertainty (stimulus contrast: *P* < 0.001, *F*_1,783_ = 48.18, ANCOVA; stimulus spread: *P* < 0.001, *F*_1,783_ = 188.72). This can be clearly seen in the stimulus uncertainty estimates, averaged across neurons (Fig. 3a). Moreover, the uncertainty manipulations did not interact significantly (*P* = 0.86, *F*_1,783_ = 0.03; Fig. 3a), suggesting that they induce stimulus uncertainty for different reasons. The average gain variability was monotonically related to the average uncertainty value (Fig. 3b). This suggests that gain variability represents the total amount of stimulus uncertainty, regardless of the source of this uncertainty. Closer examination of the behavior of individual neurons revealed that for most units, orientation uncertainty and gain variability are positively correlated (median *r* = 0.49, *P* < 0.001, Wilcoxon signed rank test; Fig. 3c).

**Figure 3.**
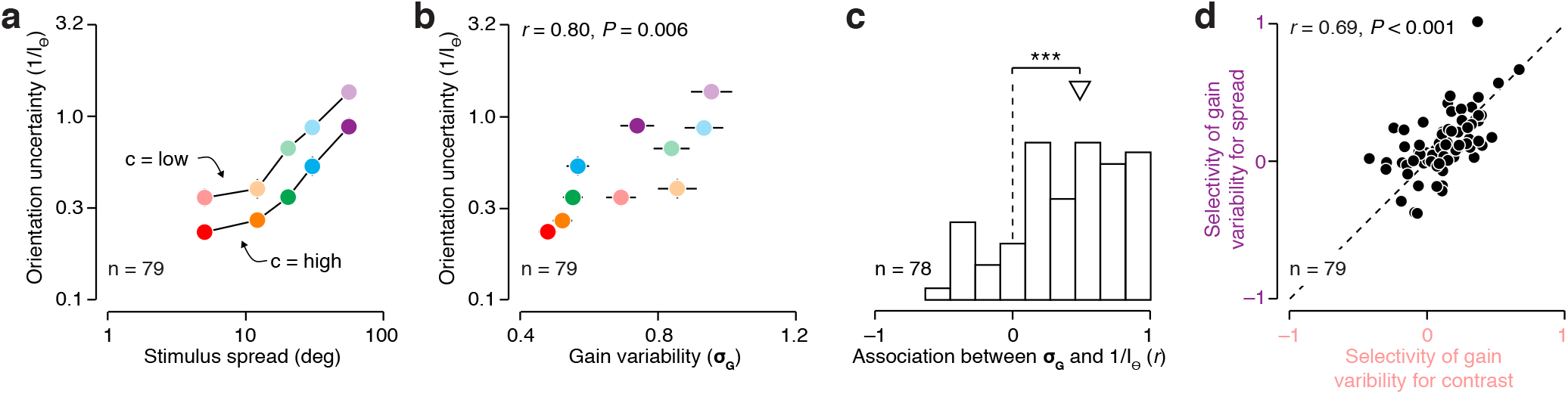
Gain variability represents stimulus uncertainty. (**a**) Orientation uncertainty as a function of stimulus contrast and spread, averaged across a population of V1 and V2 neurons. (**b**) Orientation uncertainty as a function of gain variability, averaged across a population of V1 and V2 neurons. (**c**) Distribution of the correlation between gain variability and orientation uncertainty for individual neurons. The triangle indicates the median value. (**d**) Selectivity of gain variability for stimulus spread as a function of selectivity of gain variability for stimulus contrast. Confidence intervals indicate +− 1 s.e.; *** *P* < 0.001

Finally, we asked whether the gain variability of individual neurons is tuned to stimulus uncertainty *per se*, or to a subset of the stimulus features that induce uncertainty. We singled out the most extreme stimulus manipulations, both of which induced substantial amounts of uncertainty (minimal spread at low contrast and maximal spread at high contrast). Could it be that different subsets of neurons are selective for each of these manipulations? This would question the existence of a monolithic uncertainty receptive field. We summarized each neuron’s selectivity for these manipulations by measuring the change in gain variability relative to the baseline condition (minimal spread at high contrast, see Online Methods). This statistic equals one if the stimulus manipulation increases gain variability by a factor of ten, and zero if the stimulus manipulation has no effect on gain variability (negative values indicate a decrease in gain variability). Interneuronal differences in selectivity for both manipulations were highly correlated (*r* = 0.69, *P* < 0.001; Fig. 3d). This invariance to the source of uncertainty suggests that a single mechanism could account for the uncertainty selectivity exhibited by cortical neurons.

### Representation of uncertainty across the visual hierarchy

We have, thus far, found evidence for our proposed coding scheme in the relationship between orientation uncertainty and the gain variability of orientation-selective neurons. Our theory is not limited to orientation coding, but holds that as new features are encoded along the visual hierarchy, so is their associated uncertainty. In area V2, neurons are selective for the features of visual texture, a property lacking from their V1 inputs ^21^. Our framework therefore predicts that the gain variability of V2 cells, but not V1 cells, will depend on the uncertainty about stimulus texture. To test this prediction, we analyzed responses of individual neurons in macaque V1 and V2 elicited by a set of naturalistic textures and a set of unstructured noise stimuli (Fig. 4a-b; data collected by ref. ^22^). The noise stimuli were devoid of distinctive textural features and hence induce maximal textural uncertainty – just like a uniformly dispersed stimulus would induce maximal orientation uncertainty. As predicted, noise stimuli typically elicited more gain variability than texture stimuli in V2 (median selectivity of gain variability for textural uncertainty in V2 = 0.063, *P* < 0.001; Fig. 4c-d). Neurons in V1 showed no such effect (median selectivity of gain variability in V1 = 0, *P* = 0.31; Fig. 4d). We conclude that, as neurons’ mean firing rates become selective for increasingly complex features of the visual environment, so does their gain variability for the associated uncertainty.

**Figure 4.**
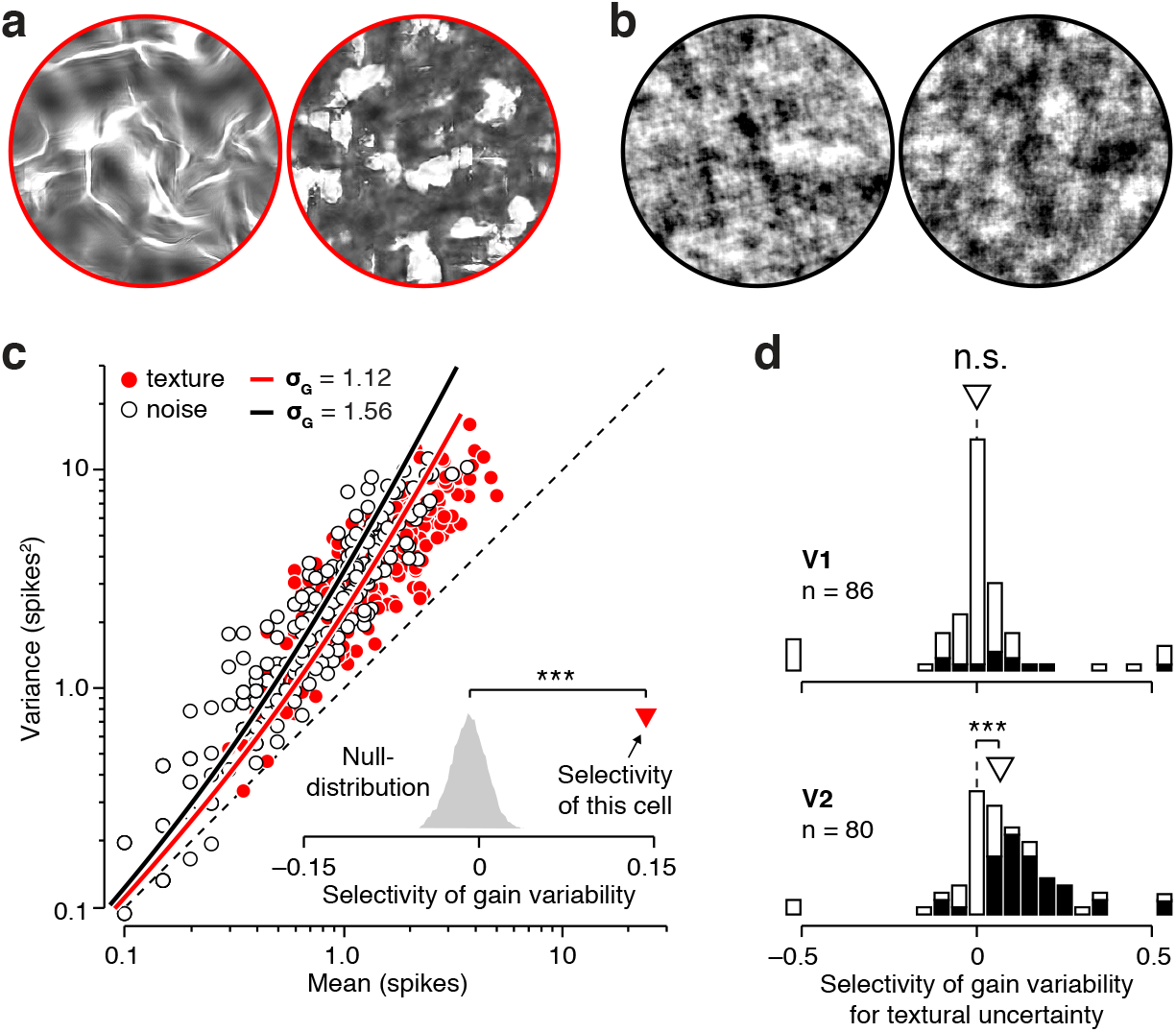
Gain variability of V2 neurons represents texture uncertainty. (**a**) Two example images with well defined textural properties (i.e. honeycomb, left; flies, right). (**b**) Noise images with ill-defined textural properties. Perceptually, the texture images are more distinct than the noise images (C.M Ziemba, J. Freeman, E.P Simoncelli, J.A Movshon, Soc Neurosci. Abstr. 573.13,2012). This differential perceptual sensitivity is reflected in the stimulus selectivity of V2 neurons, but not of V1 neurons ^21^. (**c**) Variance-to-mean relation of an example V2 cell for a set of texture images (red points), and matched noise images (white points). Lines illustrate the predictions of the modulated Poisson model, fit separately to the texture and noise stimuli. The inset shows the selectivity of gain variability for textural uncertainty expected under the null-model (grey histogram), and the selectivity estimated from the real data (red triangle). (**d**) Distribution of selectivity of gain variability for textural uncertainty for a population of V1 (top) and V2 (bottom) neurons. The dotted line illustrates the median value of the null-model. Filled bars represent cases that were significantly different from the null-model (two-sided test, *α* = 0.05). The triangle indicates the median value. n.s. not significant; *** *P* < 0.001

### The uncertainty receptive field as a consequence of stochastic divisive normalization

Which neural mechanism is general enough to support the representation of uncertainty across the visual hierarchy? Divisive normalization is a promising candidate for several reasons. First, this computation is implemented by a wide range of sensory and non-sensory circuits ^23^. Second, normalization directly controls neural response gain, and hence might also control gain variability. Finally, divisive normalization can be instantiated in image-computable models ^17,24,25^, making this a broadly testable hypothesis. We derived a stochastic formulation of the standard divisive normalization model (Fig. 5a; a related model was recently proposed in a separate context ^26^). The mean response of this model *μ* is identical to the deterministic version of the normalization model:

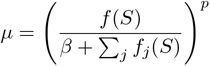

where *f* (*S*) is some function of the stimulus, *β* is a stimulus-independent constant, and *p* is a transductioxnp-eonent. The stimulus-dependent normalization factor Σ_*j*_ *f*_*j*_(*S*) reflects the aggregate activity of a large number of nearby neurons. Neural activity is noisy. We therefore make the normalization term subject to additive Gaussian noise with zero mean and variance 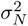. This makes the firing rate subject to stochastic gain fluctuations, and yields a simple expression for gain variability (see Online Methods):

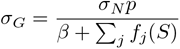

**Figure 5.**
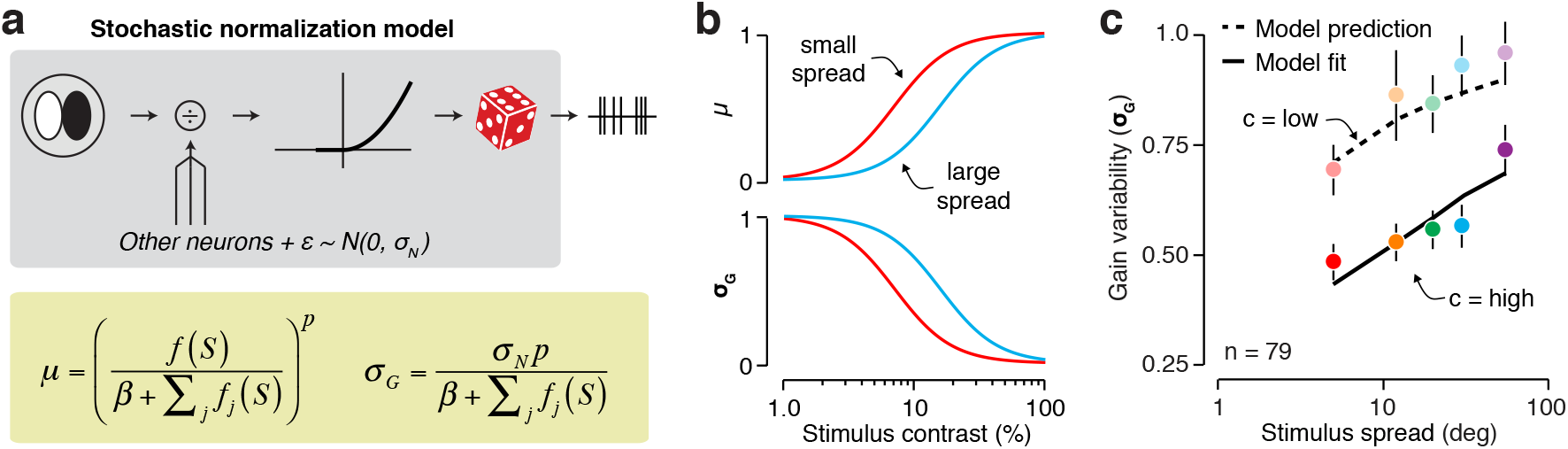
A stochastic normalization model accounts for the effects of stimulus uncertainty on gain variability. (**a**) Top: Model diagram. The response of a spatial filter is divided by the summed activity of neighboring units, an additive constant, and a noise source. The normalized signal is passed through a nonlinearity to obtain a firing rate which serves as input for a Poisson process. Bottom: equations for the mean response and gain variability under this model. (**b**) Mean response (top) and gain variability (bottom) as a function of stimulus contrast for a narrowband (red) and a broadband (blue) stimulus. (**c**) Measured gain variability compared with the prediction of the stochastic normalization model. The model was fit to average gain variability measured for high contrast stimuli (opaque points), but not to the low-contrast conditions (transparent points).

Under this stochastic normalization model, gain variability depends on the same normalization factor as the mean firing rate, and a single new parameter, the noise in the normalization signal *σ*_*N*_.

Qualitatively, this model recapitulates the trends in our data. Increasing stimulus contrast increases the normalization signal and therefore decreases gain variability (Fig. 5b). Increasing stimulus spread has the opposite effect: given a normalization pool composed of narrowly tuned neurons, the normalization signal decreases with spread, thereby increasing gain variability (Fig. 5b).

To test whether this stochastic normalization model quantitatively captures the effects of stimulus uncertainty, we fit the model to half of the data and evaluated its predictions on the other half. Specifically, we fit the only free parameter *σ*_*N*_ to the average gain variability measured for the high-contrast stimuli (all other parameters were estimated from the neurons’ mean responses, see Online Methods). This single parameter allowed the model to account for the dependency of gain variability on stimulus spread (Fig. 5c, full line; *P* = 0.17, two-sided absolute goodness-of-fit test). Keeping this free parameter constant, we predicted gain variability for the low-contrast stimulus conditions. The model correctly predicted the magnitude of the increase in gain variability (Fig. 5c, dashed line; *P* = 0.57). Hence, the uncertainty receptive field may be the functional consequence of a stochastic normalization computation.

### Gain variability exhibits slow dynamics, not fast

Our analysis suggests that gain variability in visual cortex represents stimulus uncertainty. Does this variability arise from a modulatory process with fast or slow temporal dynamics? If the process is sufficiently fast, information about stimulus uncertainty can be transmitted by individual neurons ^9^. If the process is slow, this information can only be transmitted by the joint activity of a sufficiently large population of neurons. Crucially, fast and slow modulatory processes have a different statistical signature. If the dynamics are fast, the measured variance-to-mean relation will depend on the duration of the counting window. The larger the counting window, the more within-trial variability will be averaged out, reducing the strength of measured gain fluctuations. This is not the case for a modulatory process with slow dynamics ^27^.

To address this question, we assume that stimulus-independent gain *G* is constant within temporal intervals of duration Δ*T*, but varies independently across such intervals. If this duration is longer than all measurement intervals Δ*t* (hereafter “slow” dynamics), we recover the variance-to-mean relationship described previously, which is independent of the counting window:

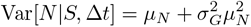

where *μ*_*N*_ = E[*N*|*S*, Δ*t*] is the mean spike count. In contrast, when Δ*T* is smaller than the shortest counting window (hereafter “fast” dynamics), the quadratic term is dampened by the counting window Δ*t*:

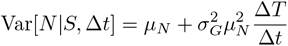

To determine whether gain fluctuations exhibit fast or slow temporal dynamics, we fit these two different versions of the modulated Poisson model to the same set of neuronal responses. We computed spike counts using differently sized counting windows (Fig. 6a), and then fit the resulting family of variance-to-mean relations imposing either fast or slow dynamics (Fig. 6b). We measured the goodness-of-fit of each model by computing its log-likelihood, and then compared both models. A recovery analysis revealed that this method distinguishes fast from slow dynamics with an accuracy of 90.15% (see Online Methods). Each unique stimulus family constitutes one point of comparison for each neuron, yielding a total of 780 data points (78 neurons × 10 stimulus families). Variance-to-mean relations were typically best described as being independent of counting window. This is evident from the responses of an example neuron. For example, notice how the fast gain dynamics model misses all the data measured with the largest counting window (Fig. 6b, right panel, blue color). The distribution of log-likelihood differences across the population supports the same conclusion (Fig. 6c; slow dynamics preferred for 89.10% of conditions, median difference = –23.4, *P* < 0.001, Wilcoxon signed rank test). In sum, gain variability seems much more likely to arise from a slow modulatory process than from a fast one.

**Figure 6.**
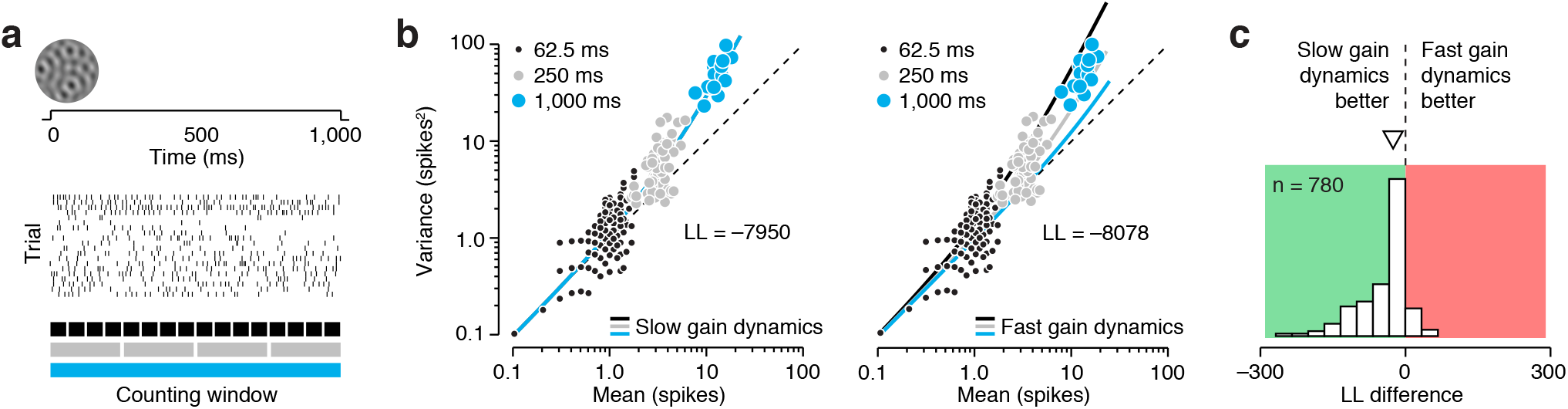
Comparison of models with fast and slow gain dynamics. (**a**) For every unique stimulus condition, we computed responses using five different spike count windows, spanning 62.5, 125, 250, 500, and 1,000 ms respectively. (**b**) Variance-to-mean relation for one stimulus family for an example V1 neuron. Responses are shown for three different counting windows: 62.5 ms (black points), 250 ms (grey points), and 1,000 ms (blue points). We fit two models to these data: one with slow gain dynamics (left panel), and one with fast gain dynamics (right panel). Slow gain dynamics predict the same variance-to-mean relationship for different counting window sizes. Fast gain dynamics predict that the variance-to-mean relationship becomes more linear with longer counting windows. We measured goodness-of-fit by computing the log likelihood of the data under each model, yielding a value of –7950 for the slow dynamics model, and of –8078 for the fast dynamics model. (**c**) Distribution of the difference in log likelihood under both models for a population of V1 and V2 neurons.

### Decoding stimulus features and uncertainty from sensory responses

Organisms have to interpret the environment almost immediately. Sensory circuits must therefore report stimulus features and their associated uncertainty on a moment-to-moment basis. Given that neuronal gain fluctuates slowly, does our proposed coding scheme enable both to be decoded quickly from sensory population activity? We investigated this using model simulations based on our experimental findings. Specifically, we simulated the activity of a population of V1 neurons that were tuned for orientation and organized in functional subpopulations (cf. cortical columns, Fig. 7a). All neurons within a column had the same stimulus selectivity *f*(*S*), resulting from linear filtering followed by divisive normalization (see Online Methods, Fig. 7b). Spikes were generated by a slowly modulated Poisson process whose gain variability was controlled by an uncertainty receptive field *u*(*S*). For simplicity, the uncertainty receptive field of all neurons had the same tuning, arising from a stochastic normalization computation (Fig. 7c). The model population thus instantiates an idealized version of the neurons we recorded from.

**Figure 7.**
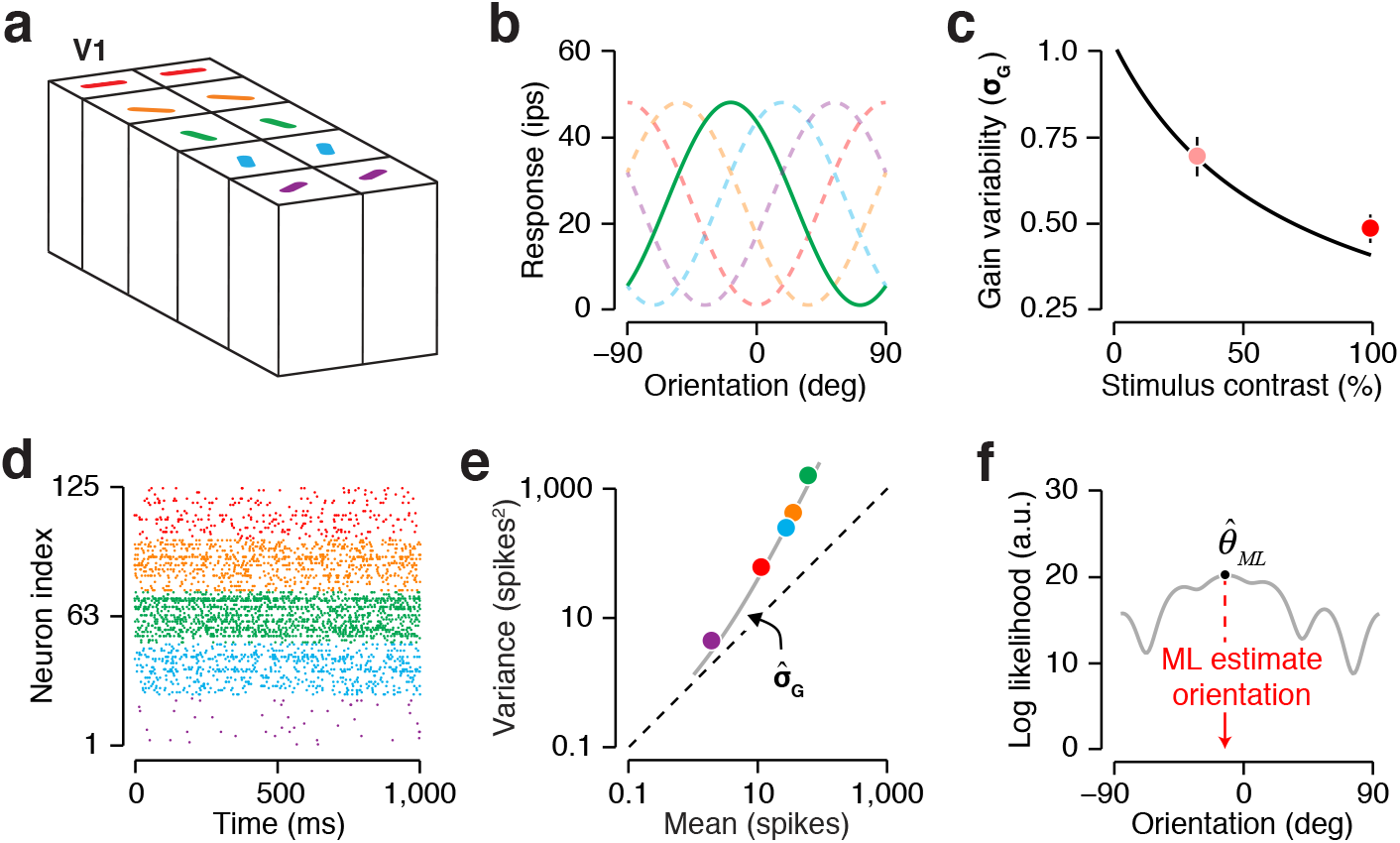
Decoding population activity. (**a**) We simulated the activity of a population of 125 V1-like neurons (5 subpopulations of 25 neurons each, loosely based on the concept of cortical columns). (**b**) All neurons within a subpopulation had the same orientation tuning function. (**c**) All neurons in the population had the same tuning of gain variability to stimulus contrast. The data points recapitulate our empirical observations. (**d**) Simulated population response for a single trial. Color indicates subpopulation. (**e**) Variance-to-mean relation for a single trial. Points summarize responses grouped by subpopulation, the line shows the predicted relation under the inferred level of gain variability. (**f**) The log-likelihood function of orientation for the same trial, given the estimated gain variability.

Consider the population response to a briefly presented sinusoidal grating (Fig. 7d). Stimulus orientation *θ* is encoded in the neurons’ average response magnitudes {*μ*_*i*_}, and stimulus uncertainty is represented by cross-neuron variability in response gain *σ*_*G*_. The optimal way to decode this information from noisy population activity is to compute the maximum likelihood estimate of both parameters. In the presence of gain fluctuations, these estimators can no longer be computed in closed form, but require an iterative estimation procedure. Given that neurons within a cortical column share the same stimulus selectivity however, super-Poisson interneuronal variance within each subpopulation can be directly attributed to gain variability. This estimate 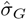 can be derived by maximum-likelihood, or using a simple, biologically plausible heuristic (Fig. 7e, see Online Methods). Equipped with this estimate of stimulus uncertainty, we can straightforwardly compute the maximum-likelihood estimate of stimulus orientation 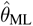 (Fig. 7f, see Online Methods). These estimates, 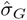 and 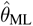, provide a useful indication of how much information regarding stimulus orientation and uncertainty is contained in the population response.

We varied stimulus orientation and uncertainty via a contrast manipulation across trials and asked how well each could be decoded from the population response on a trial-by-trial basis. For a population of 250 neurons, stimulus orientation could be decoded near perfectly when stimulus contrast was high (Fig. 8a, black symbols), but less so when contrast was low (Fig. 8a, grey symbols). This difference in performance was tracked by the simultaneously decoded gain variability. Specifically, when gain variability estimates were low, the error in orientation decoding tended to be small (Fig. 8b). But when gain variability estimates were high, the error in orientation decoding could be substantial (Fig. 8b). Gain variability estimates thus provide an instantaneously available assay of the reliability of the V1 orientation report.

**Figure 8.**
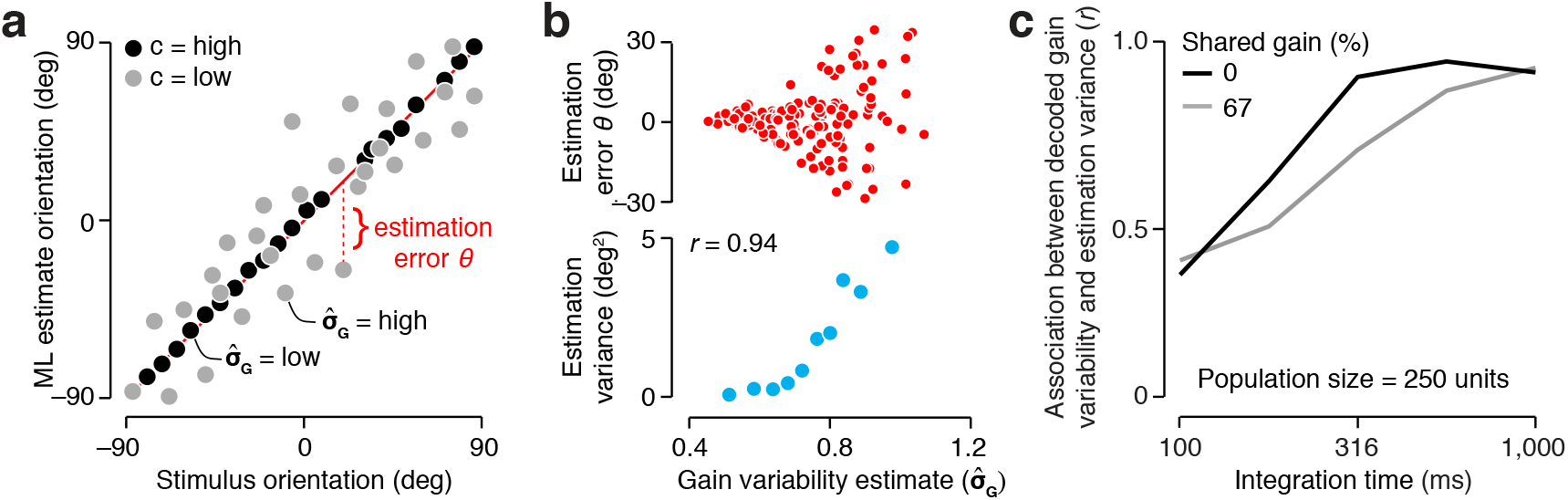
Quantitative performance of uncertainty decoding. (**a**) Estimated orientation as a function of stimulus orientation measured at high stimulus contrast (black points) and low contrast (gray points) for an example population (n = 250 neurons, integration time = 1,000 ms, shared gain fluctuations = 0%). (**b**) Top: orientation estimation error plotted against gain variability estimates. Each symbol represents a single trial. Orientation decoding error was manipulated by varying stimulus contrast across trials. Bottom: orientation estimation variance plotted against gain variability estimates. Each symbol summarizes 100 trials; trials were grouped based on their gain variability estimates (percentile 0–10, 11–20, etc.). (**c**) Effects of integration time and shared gain fluctuations on uncertainty decoding.

In the example we considered, the decoder had access to population activity realized over a one-second stimulus epoch. Moreover, all gain variability was statistically independent across neurons, in keeping with our decoder’s assumption. Decoding conditions will often be less favorable: fixations typically last only a few hundred milliseconds ^28^, and gain fluctuations can be partly shared across neurons ^15,29,30^. We wondered whether decoded gain variability would still be associated with stimulus uncertainty when read-out time is limited and gain-fluctuations are correlated. Figure 8c illustrates the evolution of this association with read-out time, split out for different levels of gain correlation. As expected, limiting read-out time and increasing gain correlations both weaken the relation between gain variability estimates and orientation uncertainty. Yet even under the most challenging conditions –read-out time = 100 ms and two thirds of gain variance is shared across neurons– the relation remained substantial (Fig. 8c). We conclude that our coding scheme enables to decode stimulus features and their uncertainty from sensory population activity under physiologically realistic conditions.

## DISCUSSION

We have introduced a new perspective on the neural code. In our view, sensory neurons report the features of the environment and the reliability of this message through two different communication channels: the mean spike count and its variance. For example, a change in stimulus orientation might alter the mean firing rate of a V1 neuron, but it will not change its gain variability. A change in orientation noise will alter the neuron’s gain variability, but need not change its mean response. We thus propose that cortical neurons behave as if two different receptive fields underlie these response statistics. While this is a new conceptualization of the neural code, we find that this behavior naturally arises from known gain control mechanisms. Gain dynamics are slow relative to behavioral time-scales, hence gain variability cannot be communicated quickly by individual neurons. Nevertheless, we have shown through model simulations that our coding scheme enables sensory populations to rapidly report stimulus features and their uncertainty to downstream circuits, even when gain variability is highly correlated across neurons.

Our view of the representation of uncertainty in cortex sits in between existing theories. Probabilistic population codes (PPC) postulate that the first volley of feedforward spikes implicitly encodes a parametric distribution over stimulus features ^7^. We share this view and extend it by proposing that not just the mean activity but also inter-neuronal variance constrain the parameters of this distribution. The simplicity of parametric, feedforward encoding makes both schemes amenable to straightforward decoding. The addition of a separate uncertainty channel makes our representation richer than traditional PPC, allowing the integration of uncertainty stemming from sources that don’t change the mean response into the representation. There is some evidence that sensory systems exploit this extra bandwidth. For example, when an observer pays attention to a visual stimulus, perceptual uncertainty can be greatly reduced ^31^. In early visual cortex, this behavioral effect is associated with a mild increase in mean response ^32^, and a comparatively strong reduction in response variability ^14^. Moreover, visual attention appears to achieve these effects by employing sensory normalization mechanisms ^33,34^, and specifically reduces neural gain variability ^30,35^.

Our interpretation of response variance bears similarity to the other major theory of uncertainty representation, which holds that neural activity represents a sample from the posterior distribution over stimulus features ^9,36^. However, we differ by stating that neural activity does not represent an arbitrarily flexible posterior distribution, but rather the peak and width of a simple distribution determined by the classical and uncertainty receptive fields. This conceptual simplicity allowed us to straightforwardly fit the uncertainty receptive field to V1 spiking data (Fig. 5c). The notion of a population-based representation of uncertainty also differentiates our view from time-based sampling models ^9,36^. Although our model explains the structure of sensory response variability found during passive viewing, behavioral requirements may induce additional task-specific variability ^37–39^, and sampling models have explained that component of neural response variability ^36^. A complete account of neural uncertainty representation might thus require combining our feedforward coding scheme with a recurrent, sampling-based one –an approach that has been shown to combine the advantages of both in machine systems ^40^.

To test our theory, we relied on stimulus manipulations that impair perceptual orientation judgments, and we verified that they reduced the coding capacity of orientation-selective neurons (Fig. 2c, Fig. 3a). This approach can directly be extended to other stimulus features, visual areas, and sensory systems to investigate the generality of the uncertainty receptive field. As a first step, we have shown that V2 cells, whose mean firing rate is selective for textural properties ^21^, modulate their gain variability according to uncertainty in visual texture. Crucially, V1 cells, which lack this selectivity, also fail to report this uncertainty. This suggests that, along a sensory processing cascade, selectivity for novel stimulus features and an assessment of their reliability jointly emerge. Why is this so? The sensory neurons that are the first in the hierarchy to represent a particular feature are uniquely positioned to judge the quality of the evidence for that feature. Downstream areas can inherit the feature report, but neural stochasticity entails that uncertainty about this feature can only grow along the hierarchy. Consistent with this, visual areas downstream of V1 exhibit orientation selectivity, but this selectivity is accompanied by systematically increasing levels of gain variability ^15^.

Our results offer a novel view of the structural organization of sensory cortex. Its columnar organization has been known for many decades ^16,41,42^, yet the computational benefit of this structure has remained elusive ^43^. In our coding scheme, estimating inter-neuronal gain variability is greatly facilitated by the presence of sub-populations of sensory neurons that share the same stimulus selectivity. In particular, this allows a decoder to infer stimulus uncertainty without detailed knowledge of the sensory neurons’ classical receptive field. Whether downstream circuits actually employ this read-out scheme can only be ascertained from an awake, behaving paradigm that requires taking stimulus uncertainty into account. A recent study of this kind found that orientation uncertainty represented by V1 populations (estimated using a flexible, model-agnostic approach) does indeed inform animals’ choice behavior ^44^. We believe that this paradigm can be leveraged to test our and other theories, and ultimately will uncover which aspect of neural activity informs perceptual uncertainty estimates.

Finally, our results reveal a strong connection between biological and machine inference under uncertainty. Recent years have witnessed the development of a new class of highly scalable artificial inference methods ^45,46^. Like our coding scheme, these methods forfeit exact inference which often requires costly iterative procedures ^47^ in favor of simple, parametric distributions that can be computed in a feedforward manner. The resulting efficiency and scalability has enabled solving highly complex real-world problems such as scene understanding ^48^, autonomous driving ^49^, and robotic manipulation ^50^. Biological systems face similarly complex tasks and environments, and may therefore also have opted for inference methods that are simple and powerful.

## METHODS

### Physiology

The data analyzed here were previously published, and the full methods are provided there (see ref. ^17^ for the orientation experiment, and ref. ^22^ for the texture experiment). In brief, all recordings were made from anesthetized, paralyzed, adult macaque monkeys. Surgical preparation methods are reported in detail in ref. ^51^. Anesthesia was maintained with infusion of sufentanil citrate (6-30 g kg^−1^h^−1^) and paralysis with infusion of vecuronium bromide (Norcuron; 0.1 mg kg^−1^h^−1^) in isotonic dextrose-Normosol solution. All experiments were conducted in compliance with the NIH’s *Guide for the Care and Use of Laboratory Animals*, and with approval of the New York University Animal welfare committee. Extracellular recordings from individual neurons were made with quartz-platinum-tungsten microelectrodes (Thomas Recording), advanced mechanically into the brain through a craniotomy and small durotomy. V1 was distinguished from V2 on the basis of depth from the cortical surface and changes in the receptive field location of the recorded units.

### Visual Stimulation

In the orientation experiment, stimuli consisted of Gaussian orientation mixtures, created by summing nine sinusoidal gratings whose orientations were spaced at 20° intervals and whose orientation-dependent contrasts followed a circular Gaussian profile centered on a particular orientation (spread 0–55°). The drift rate of each stimulus component was selected at random from a Gaussian distribution centered on the preferred rate, with an SD equal to 1/5 this value, resulting in an incoherently drifting mixture. In total, ten stimulus families (five spread levels × two contrast levels) were presented at 16 different orientations.

In the texture experiment, stimuli were generated using the texture analysis-synthesis procedure introduced by Portilla and Simoncelli ^52^. Fifteen different grayscale photographs of visual texture served as prototypes. From each of these source images, two times 15 samples were synthesized (one set of “naturalistic textures”, and one set of “unstructured noise stimuli”). The synthetic textures preserved the second-order statistics as well as some of the higher-order statistics of the source images; the noise stimuli only preserved the second-order statistics ^22^.

In both experiments, stimuli were presented in random order for either 1,000 ms (orientation experiment) or 100 ms (texture experiment), and typically repeated 10 times (orientation experiment) or 20 times (texture experiment).

### Data Analysis

For all analyses of the orientation experiment but one, we counted spikes within a 1,000 ms window following response onset. One analysis sought to compare spiking models with slow vs fast gain dynamics (fig. 6). Here, we used five different counting windows (62.5, 125, 250, 500, and 1,000 ms). For the analysis of the texture experiment, we computed spike counts using a 100 ms window aligned to the response onset.

### Quantifying neural stimulus uncertainty

Using standard tools from information theory ^20^, we quantified neural stimulus uncertainty in the orientation domain as the inverse of a neuron’s Fisher Information for a given stimulus family. If neural responses arise from a Poisson process, this statistic can be simply written as a function of the measured tuning curve, *h*(*θ*):

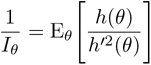

where *h*′(*θ*) is the derivative of the tuning curve ^53^. This statistic has the benefit that its value only depends on the measured mean responses, and is independent of the level of gain fluctuations. Associations between gain variability and stimulus uncertainty (Fig. 2f, Fig. 3b,c) can thus not arise for trivial reasons. This is not true of alternative estimators of uncertainty which rely on empirical measurements of response variance rather than a Poisson assumption.

### Measuring gain variability

We measured gain variability using the method introduced by ref. ^15^. Specifically, we described responses of individual neurons with a model in which spikes are generated by a Poisson process whose rate is the product of a stimulus dependent drive and a stimulus independent gain. We assumed that gain is constant within a trial and distributed across trials according to a gamma distribution with mean 1 and variance 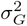. We esti-mated this parameter by maximizing the likelihood of the full set of observed spike counts for a given stimulus family under a negative binomial distribution ^15^ (Fig. 2d, Fig. 4c).

We computed the selectivity of gain variability for induced stimulus uncertainty (Fig. 3d, Fig. 4d) by taking the common logarithm of the ratio of two *σ*_*G*_ estimates: one measured in the presence of the uncertainty-inducing manipulation (numerator), and one measured in its absence (denominator). For the texture experiment, we performed a significance test on this statistic (Fig. 4c, inset). For each neuron, we obtained a null-distribution by first estimating gain variability from the combination of all stimulus conditions. Next, we used this value and the empirically observed mean responses to simulate 100 synthetic data-sets. We then separately estimated gain variability for the subset of synthetic responses to texture and noise stimuli, respectively. We used these values to compute the expected distribution of the selectivity-index if there were no underlying difference in gain variability between texture and noise stimuli. We deem the empirically obtained selectivity value significant if it falls outside of the central 95 percent interval of this distribution (estimated from 10,000 samples per neuron).

### Analysis of gain dynamics

We sought to determine whether neural gain fluctuations are better described as having fast or slow dynamics. For a slow modulatory process, the variance-to-mean relationship is independent of counting window; for a fast process, this relation changes in a predictable manner with window size (see equations in Results). To leverage this insight, we counted the same set of spikes with windows of different duration, and then fit both a fast- and a slow-dynamics model to the resulting data-set. The largest counting window (1,000 ms) contributes one observation per trial; the smallest window (62.5 ms) contributes sixteen observations per trial. To determine the log-likelihood of the models for an entire data-set, we treat all observations as being statistically independent. This is not strictly correct, as each spike is counted multiple times (exactly once per window size). To assess the effectiveness of our model comparison procedure, we performed a recovery analysis. For each measured variance-to-mean relation (one per neuron per stimulus family), we synthesized 1,000 data-sets imposing slow gain dynamics, and 1,000 data-sets imposing fast gain dynamics. The generating parameters were the empirically observed mean counts as measured with a 62.5 ms window, and the gain variability estimate obtained under a 1,000 ms window. We then fit the slow- and fast-dynamics model to each synthetic data-set, and compared their goodness-of-fit in exactly the same manner as we did for the real data. When the ground-truth was slow dynamics, the slow dynamics model was preferred in 99.51% of cases; when the ground-truth was fast dynamics, the fast dynamics model was preferred in 80.78% of cases. We deem our method to be fairly sensitive, and slightly biased in favor of slow dynamics. If slow and fast dynamics were equally probable in the population, our method would identify the slow dynamics model as the winner in 59.37% of cases. This number is larger than 50%, but much smaller than the empirical outcome (slow dynamics favored in 89.10% of cases).

### Fitting the stochastic normalization model

The canonical divisive normalization model describes the firing rate *λ*_*i*_ of a neuron *i* in response to a stimulus *S* as some function of the stimulus drive *f*_*i*_(*S*) divided by the sum of stimulus-dependent drive to *j* neighboring neurons Σ_*j*_ *f*_*j*_(*S*) and a stimulus-independent constant *β*, with transduction exponent *p*:

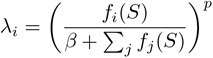

Because neighboring neurons are stochastic, we modeled the aggregate stochasticity of the normalization pool with stimulus-independent additive Gaussian noise 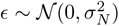:

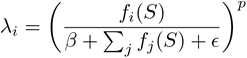

If the magnitude of the noise *ϵ* is sufficiently small, we can use a Taylor expansion to obtain the mean and standard deviation of the firing rate:

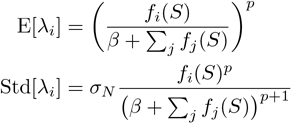

Equating these expressions to those obtained from the modulated Poisson model (recall E[*λ*_*i*_] = *f* (*S*), Std[*λ*_*i*_] = *f* (*S*)*σ*_*G*_) results in an expression of gain variability:

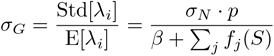

Although the noise term *σ*_*N*_ is stimulus-independent, divisive normalization causes gain variability to depend on the stimulus through the denominator of this expression. We investigated the adequacy of this equation by fitting the stochastic normalization model to the average gain variability measured for the high-contrast mixture stimuli. We opted to constrain the model as much as possible. Rather than fitting the transduction exponent *p* and the stimulus-independent normalization constant *β* to these data, we used the average estimates of both parameters obtained by fitting the neurons’ mean responses with a divisive normalization model ^17^ (*p* = 2, *β* = 0.64). The stimulus-dependent normalization Σ_*j*_ *f*_*j*_(*S*) was computed by simulating responses of a fixed pool of neurons with a diverse set of tuning properties ^17^. The final free parameter, *σ*_*N*_, was estimated by minimizing the sum of squared error between predicted and observed *σ*_*G*_ (Fig. 5c, full line).

### Decoding orientation and its uncertainty

Stimulus uncertainty and orientation were decoded on a trial-by-trial basis from a simulated population of 250 V1-like neurons whose tuning curves evenly tiled the orientation domain (*n* = 10 subpopulations of *m* = 25 units each). Within a subpopulation *i*, all neurons *j* had the same orientation selectivity *f*_*i*_(S). All neurons had the same uncertainty receptive field whose shape was determined by parameters fit to neural data (*σ*_*N*_ = 0.34, *p* = 2, *β* = 0.64), resulting in a single value *σ*_*G*_ for the gain variability of the entire population. We decode orientation *S* and uncertainty *σ*_*G*_ by first evaluating their likelihood, given a collection of *n* * *m* spike counts 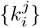 afropmopulation of independent, modulated Poisson neurons with gamma-distributed gain:

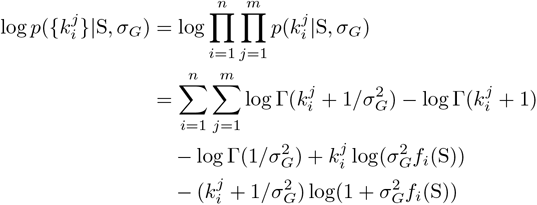

Although the maximum likelihood estimators of S and *σ*_*G*_ are not available in closed form, they can be obtained using an iterative, gradient-based procedure. This procedure can be simplified given that neurons within a column share the same firing rate, which can be estimated by averaging over these neurons:

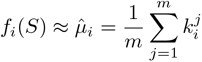

with the approximation becoming exact in the limit of a large sub-population size *m*. Finding the optimal value of the gain variability then reduces to a one-dimensional problem:

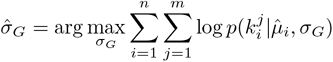

This is the decoded gain variability reported in Fig. 8. However, its expression can be further simplified i nto a heuristic estimator by noticing that, because neurons within a sub-population are identically tuned, inter-neuronal variance can be directly attributed to gain variability:

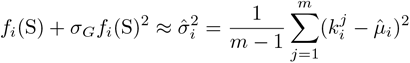

with the approximation becoming exact in the limit of a large sub-population size *m*. This leaves us with a simple estimator 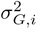 for the gain variability within a sub-population, wcahnicbhe straightforwardly extended to the entire population by summing statics across sub-populations:

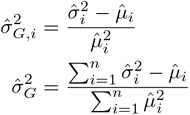

For the conditions reported in Fig. 8, this heuristic estimator closely approximates the maximum-likelihood estimate.

Equipped with an estimate 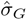 of the gain variability, computing the maximum-likelihood estimate of orientation reduces to another one-dimensional problem:

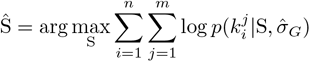

To assess the quality of uncertainty and orientation decoding, we measured the orientation decoding error on a trial-by-trial basis (Fig. 8a). Each simulation included 5 repetitions of 20 different contrasts and 10 different orientations, yielding a total of 1000 trials. We sorted and binned the trials according to the estimated gain variability 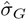. Within each bin, we computed the mean squared error across trials, and compared it to the average gain variability estimate of that bin (Fig. 8b). The reported association between these two quantities (Fig. 8c), is their Pearson correlation, appropriately averaged across 100 repeats of the simulation.

To assess the effect of interneuronal gain correlations, we varied the amount of gain correlation while keeping the total amount of gain variability constant. Specifically, we created two gain variables *G*_*s*_ and *G*_*p*_ that were shared and private respectively, both of which had unit mean and a variance equal to 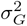. Each neuron was modulated by its own gain *G* = *λG*_*s*_ + (1 − *λ*)*G*_*p*_ where *λ* ∈ [0, 1]. When *λ* = 0, all gain variability is statistically independent across the population, when *λ* > 0, interneuronal gain fluctuations are positively correlated. We chose *λ* ∈ {0, 0.67} to span a physiologically plausible range ^15, 54^.

